# Stratifying major depressive disorder by polygenic risk for schizophrenia in relation to structural brain measures

**DOI:** 10.1101/663724

**Authors:** Mathew A. Harris, Xueyi Shen, Simon R. Cox, Jude Gibson, Mark J. Adams, Toni-Kim Clarke, Ian J. Deary, Stephen M. Lawrie, Andrew M. McIntosh, Heather C. Whalley

## Abstract

**Background:** Substantial clinical heterogeneity of major depressive disorder (MDD) suggests it may group together individuals with diverse aetiologies. Identifying distinct subtypes should lead to more effective diagnosis and treatment, while providing more useful targets for further research. Genetic and clinical overlap between MDD and schizophrenia (SCZ) suggests an MDD subtype may share underlying mechanisms with SCZ.

**Methods:** The present study investigated whether a neurobiologically distinct subtype of MDD could be identified by SCZ polygenic risk score (PRS). We explored interactive effects between SCZ PRS and MDD case/control status on a range of cortical, subcortical and white matter metrics among 2,370 male and 2,574 female UK Biobank participants.

**Results:** There was a significant SCZ PRS by MDD interaction for rostral anterior cingulate cortex (RACC) thickness (β=.191, q=.043). This was driven by a *positive* association between SCZ PRS and RACC thickness among MDD cases (β=.098, p=.026), compared to a negative association among controls (β=–.087, p=.002). MDD cases with low SCZ PRS showed thinner RACC, although the opposite difference for high-SCZ-PRS cases was not significant. There were nominal interactions for other brain metrics, but none remained significant after correcting for multiple comparisons.

**Conclusions:** Our significant results indicate that MDD case-control differences in RACC thickness vary as a function of SCZ PRS. Although this was not the case for most other brain measures assessed, our specific findings still provide some further evidence that MDD in the presence of high genetic risk for SCZ is subtly neurobiologically distinct from MDD in general.

## Introduction

Major depressive disorder (MDD) is a prevalent and frequently disabling psychiatric disorder, associated with prolonged low mood, although many unique symptom combinations may lead to the same diagnosis (Gallo & Rabbins, 1999; Kennedy, 2008). The high clinical heterogeneity suggests that MDD may group together a number of disease subtypes (Fava et al., 1997; Baumesiter & Parker, 2012; van Loo et al., 2014). Differences in clinical presentation may reflect differences in aetiology and underlying neurobiological mechanisms. Identifying subtypes of MDD is therefore likely to be a crucial step toward more effective diagnosis and treatment of affected individuals, as well as providing better targets for genomic and neurobiological studies.

The heritability of MDD is estimated to be about 37% (Sullivan et al., 2000), determined by a large number of alleles each of small effect (Ripke et al., 2013). MDD subtypes may differ in terms of which alleles (Flint & Kendler, 2014) and which biological mechanisms (Hasler et al., 2004) are involved. For example, Milaneschi et al. (2017) demonstrated only a very small genetic overlap between typical MDD (associated with decreased appetite and weight) and an atypical subtype (associated with increased appetite and weight). Focusing more specifically on stratifying MDD by genetic factors, Howard et al. (2017) recently identified 10 variables related to genetic subgroups that might partly account for the heterogeneity of MDD. Further investigation of specific genetic subgroups may help to identify more clinically and empirically useful MDD subtypes.

Some genes that contribute to MDD overlap with those underlying schizophrenia (SCZ; Schulze et al., 2014) and the two disorders have a genetic correlation of r_g_=.43 (Cross-Disorder Group of the Psychiatric Genetics Consortium et al., 2013). Individuals with MDD also at high polygenic risk of SCZ have been previously shown to differ in terms of clinical and behavioural phenotypes. Whalley et al. (2016) observed higher neuroticism and psychological distress among controls with a higher SCZ polygenic risk score (PRS), but not among high-SCZ-PRS MDD cases. Additionally, Power et al. (2017) found that MDD subtypes based on age of onset differed in terms of SCZ PRS, and early-onset cases have been found to show greater deficits in frontal (Jaworska et al., 2014) and limbic (MacMaster et al., 2008; Clark et al., 2018) regions. This may suggest a different aetiology and perhaps different neurobiological profile of MDD in the presence of high genetic risk for SCZ, i.e. an SCZ-risk-related MDD subtype. It seems that some of the genes that may contribute to both SCZ and MDD are involved in regulating synaptic function and the excitability of prefrontal neurons (Howard et al., 2019). However, whether subtypes of MDD based on genetic loading for SCZ are distinct in terms of brain structure has yet to be determined.

MDD is typically associated with fronto-limbic deficits, including reduced cortical volume/thickness in prefrontal regions (Salvadore et al., 2011; Rodríguez-Cano et al., 2014; Schmaal et al., 2017), and smaller subcortical limbic structures (Kim et al., 2008; Lu et al., 2016; Schmaal et al., 2016). Reductions in integrity of connecting white matter tracts have also been observed (Cole et al., 2012; Bessette et al., 2014; Shen et al., 2017). Fronto-limbic impairments are also apparent in SCZ (Ross et al., 2006; Keshavan et al., 2008; Yao et al., 2013), along with more widespread white matter deficits (Ellison-Wright & Bullmore, 2009) and greater cortical reductions in temporal regions (Wong & van Tol, 2003). Furthermore, some SCZ-like neuroimaging markers, such as reduced hippocampal, thalamic and prefrontal volumes, appear in individuals who are genetically predisposed to SCZ but remain healthy (Lawrie et al., 2001; Boos et al., 2007). As of yet, the influences of SCZ risk on such neurobiological measures in the presence of MDD are unclear.

Following on from previous work stratifying MDD traits by PRS for SCZ (Whalley et al., 2016), we aimed to measure the neuroimaging associations of MDD as a function of SCZ PRS, testing for further evidence of MDD subtype heterogeneity related to SCZ risk. We assessed interactive effects between SCZ PRS and MDD case/control status on measures of cortical, subcortical and white matter structure. Based on previous evidence of neuroimaging abnormalities in healthy subjects at high genetic risk for SCZ (Lawrie et al., 2001; Boos et al., 2007), we expected negative correlations between SCZ PRS and brain measures among both cases and controls, with significant interactions between MDD and SCZ PRS indicating a specific subtype. Without any specific hypotheses regarding the neurobiological bases of differences in neuroticism and distress among high-SCZ-risk MDD cases, we took an exploratory approach, testing SCZ PRS by MDD interactions for structural measures covering the entire brain. We did, however, anticipate interactions in areas that normally show deficits in MDD, such as frontal cortical regions, limbic structures and connecting white matter tracts. Any significant interactions, indicating that MDD’s associations with neuroimaging measures differed as a function of SCZ PRS, would provide evidence of an SCZ-risk-related MDD subtype.

## Methods

### Participants

Data were derived from 4,944 participants in the large-scale prospective epidemiological study, UK Biobank (http://www/ukbiobank.ac.uk). UK Biobank includes data on 503,325 members of the general UK population, recruited between 2006 and 2010 (Allen et al., 2012). Participants originally provided information on a wide range of health, lifestyle, environment and other variables, and the majority have also provided genetic data. The project aims to submit approximately 100,000 of the participants to a full-body imaging protocol, which includes the acquisition of neuroimaging data in several modalities (Miller et al., 2016). The present study focused on 4,944 UK Biobank participants for whom both genetic and brain imaging data have already been acquired, who also had data on MDD and relevant covariates, described below. Participants who reported a diagnosis of schizophrenia were excluded from all analyses. Descriptive statistics for included participants are reported in Table 1.

**Table 1.** Descriptive statistics for MDD cases and controls, and group differences.

|  | Cases | Controls | Difference |
| --- | --- | --- | --- |
| Subjects (N, %) | 1674 (33.86) | 3270 (66.14) | $\chi^2_1=1029.10$, $p<.001$ *** |
| Age (years; M, SD) | 61.23 (7.20) | 63.69 (7.21) | $t_{3249}=-11.15$, $p<.001$ *** |
| Sex (N, %) |  |  |  |
| Males | 625 (37.34) | 1745 (53.36) | $\chi^2_1=113.33$, $p<.001$ *** |
| Females | 1049 (62.66) | 1525 (46.64) |  |
| WBV (cm<sup>3</sup>; M, SD) | 1194.91 (111.77) | 1212.24 (114.99) | $t_{3458}=-5.11$, $p<.001$ *** |
| SCZ PRS (z-score; M, SD) |  |  |  |
| Threshold .01 | 0.03 (0.98) | -0.03 (1.01) | $t_{3457}=1.87$, $p=.06$ |
| Threshold .05 | 0.02 (0.98) | -0.03 (1.01) | $t_{3476}=1.91$, $p=.06$ |
| Threshold .10 | 0.03 (0.98) | -0.04 (1.02) | $t_{3482}=2.28$, $p=.02$ * |
| Threshold .50 | 0.03 (0.99) | -0.05 (1.01) | $t_{3430}=2.91$, $p<.01$ ** |
| All SNPs | 0.04 (0.99) | -0.05 (1.01) | $t_{3428}=3.01$, $p<.01$ ** |
| Imaging data (N, %) |  |  |  |
| Cortical metrics | 640 (38.23) | 1237 (37.83) | $\chi^2_1=0.09$, $p=.77$ |
| (replication sample) | 794 (49.50) | 1604 (51.19) | $\chi^2_1=1.16$, $p=.28$ |
| Subcortical volumes | 1674 (100.00) | 3270 (100.00) | $\chi^2_1=0.00$, $p>.99$ |
| DTI measures | 1474 (88.05) | 2946 (90.09) | $\chi^2_1=0.55$, $p=.79$ |

*Notes.* MDD = major depressive disorder; WBV = whole brain volume; SCZ = schizophrenia; PRS = polygenic risk score; DTI = diffusion tensor imaging. Group differences were assessed using $\chi^2$ tests and two-sample t-tests. \*, \*\* and \*\*\* represent significant results at $p<.05$ , $p<.01$ and $p<.001$ , respectively.

### Polygenic risk for schizophrenia

UK Biobank blood samples were genotyped using either the UK BiLEVE array or UK Biobank Axiom array, and quality controlled at the Wellcome Trust Centre for Human Genetics (Wain et al., 2015; Hagenaars et al., 2016). Further quality control measures included removal of participants based on missingness, relatedness, non-British ancestry and gender mismatch. Subsequent processing involved removal of SNPs with minor allele frequency <1%, and clump-based linkage disequilibrium pruning (r^2^<.25 within a 200bp window). PRSs for SCZ were calculated for each participant using PRSice (Eusden et al., 2015) and genome-wide association study summary data (Schizophrenia Working Group of the Psychiatric Genetics Consortium, 2014), including SNPs selected according to the significance of their associations with the phenotype at thresholds of p<.01, p<.05, p<.10, p<.50 and all SNPs.

### Major depressive disorder

As part of a web-based mental health questionnaire, UK Biobank participants completed the short form of the Composite International Diagnostic Interview (CIDI-SF; Kessler et al., 1998). Section A of the CIDI-SF focuses on MDD symptomatology, providing a score of between zero and eight. Participants with a score of five or more were classified as lifelong MDD cases; those with a score of four or less as controls. Participants who scored highly but did not report having experienced core symptoms of either low mood or anhedonia were excluded. CIDI-SF controls who had previously reported a past diagnosis of depression or showed signs of MDD on the 9-item Patient Health Questionnaire (PHQ-9; Kroenke et al., 2001) were also excluded (N=286). The resulting CIDI-assessed measure of MDD therefore included 1,674 cases and 3,270 controls (Table 1). Some of the earlier scanned subjects completed the web-based questionnaire up to 2.7 years later, but most completed it within a year of imaging data acquisition.

### Brain imaging data

Full information on imaging data acquisition and processing are provided in supplementary materials and previous publications (Miller et al., 2016; Alfaro-Almagro et al., 2018). To summarise, all structural and diffusion data were acquired on the same 3T scanner using the same protocol. Cortical metrics for 27 regions (N=1,877) were derived locally using FreeSurfer version 5.3 (Dale et al., 1999; Fischl et al., 1999; Fischl et al., 2004; Desikan et al., 2006). Volumes of seven subcortical structures (N=4,944) were derived by the UK Biobank imaging team using FIRST (Patenaude et al., 2011). Fractional anisotropy (FA) and mean diffusivity (MD) of 15 white matter tracts (N=4,420) were also derived by the UK Biobank imaging team using bedpostx and probtrackx (Behrens et al., 2007), and then AutoPtx (de Groot et al., 2013).

At the time of beginning this study, subcortical volumes and white matter metrics were available for the first two releases of UK Biobank imaging data, while cortical metrics were available for only the first release. Following initial submission of our results for publication, cortical metrics became available for the second release as well, including for 794 MDD cases and 1,604 controls who met the criteria for inclusion in this study. We have therefore used these additional data to replicate any significant findings discovered in the first release.

### Additional covariates

In relation to SCZ PRS, genotyping array and the first 15 ancestry principal components (controlling for population stratification within the sample) were included as covariates in all analyses. We used as many as 15 principal components to control for the substantial genetic clustering recently found in UK Biobank (Abdellaoui et al., 2018). Including more than 15 made negligible difference to our results. In relation to neuroimaging data, x, y and z coordinates of head position within the MRI scanner were included as covariates, along with a measure of whole-brain volume (including grey matter, white matter and ventricles), derived from the UK Biobank image processing pipeline (Alfaro-Almagro et al., 2018). Additionally, age in years at time of scanning, age^2^ and sex were included in all analyses.

### Statistical analysis

Data were analysed in R version 3.2.3 (R Core Team, 2013). As additional quality control, outliers, defined as further than three standard deviations from the mean, were removed from all neuroimaging measures. Left and right measures were then combined bilaterally, as above; a general lack of hemispheric effects in the same UK Biobank subcortical and white matter phenotypes has been reported previously (Shen et al., 2017). We performed analyses of case-control differences in group size, sex and SCZ PRS, reported in Table 1. For each regional brain measure and each PRS threshold, we then ran linear regression models to estimate the strength of the interactive effect between SCZ PRS and MDD in all participants, controlling for the main effects of each, as well as effects of age, age^2^, genotyping array, the first 15 principal components of genetic ancestry, whole-brain volume and scanner head position coordinates. FDR correction was applied to p values across all interaction effects together.

As main effects of SCZ PRS among MDD cases would not demonstrate whether the effects were specific to cases, only interactions between MDD and SCZ PRS could provide indications of aetiological differences between MDD cases at high and low SCZ risk. However, where interactions were significant, we assessed main effects of SCZ PRS among MDD cases and controls separately, and tested case-control differences among those above and below mean SCZ PRS. Together, these analyses provided further information on how MDD and SCZ risk interact. We additionally tested whether significant interactions were influenced by a subgroup of MDD cases with both repeated k-mean and hierarchical cluster analyses, using package “fpc” (Henning, 2015) in addition to base packages. Further packages “ggplot2” (Wickham & Chang, 2016) and “ellipse” (Murdoch & Chow, 2013) were also used to create figures. Results are reported in terms of standardised β coefficients, and p values and q values (FDR-corrected p values) <.05 are considered significant.

## Results

Descriptive statistics are reported for MDD cases and controls in Table 1. MDD cases were younger than controls (t_3252_=-11.15, p<.001), a higher proportion of cases were female (χ^2^_1_=113.33, p<.001), and WBV was lower among cases (t_3460_=5.11, p<.001). These variables were controlled in all subsequent analyses. Mean PRS was slightly higher among cases at thresholds. 10 (t_3484_=2.28, p=.02), .50 (t_3430_=2.91, p<.01) and all SNPs (t_3427_=3.01, p<.01). All SCZ PRS by MDD interactions tested for both MDD measures are summarised in Figure 1; within each measure, fronto-limbic areas are presented first.

**Figure 1.**
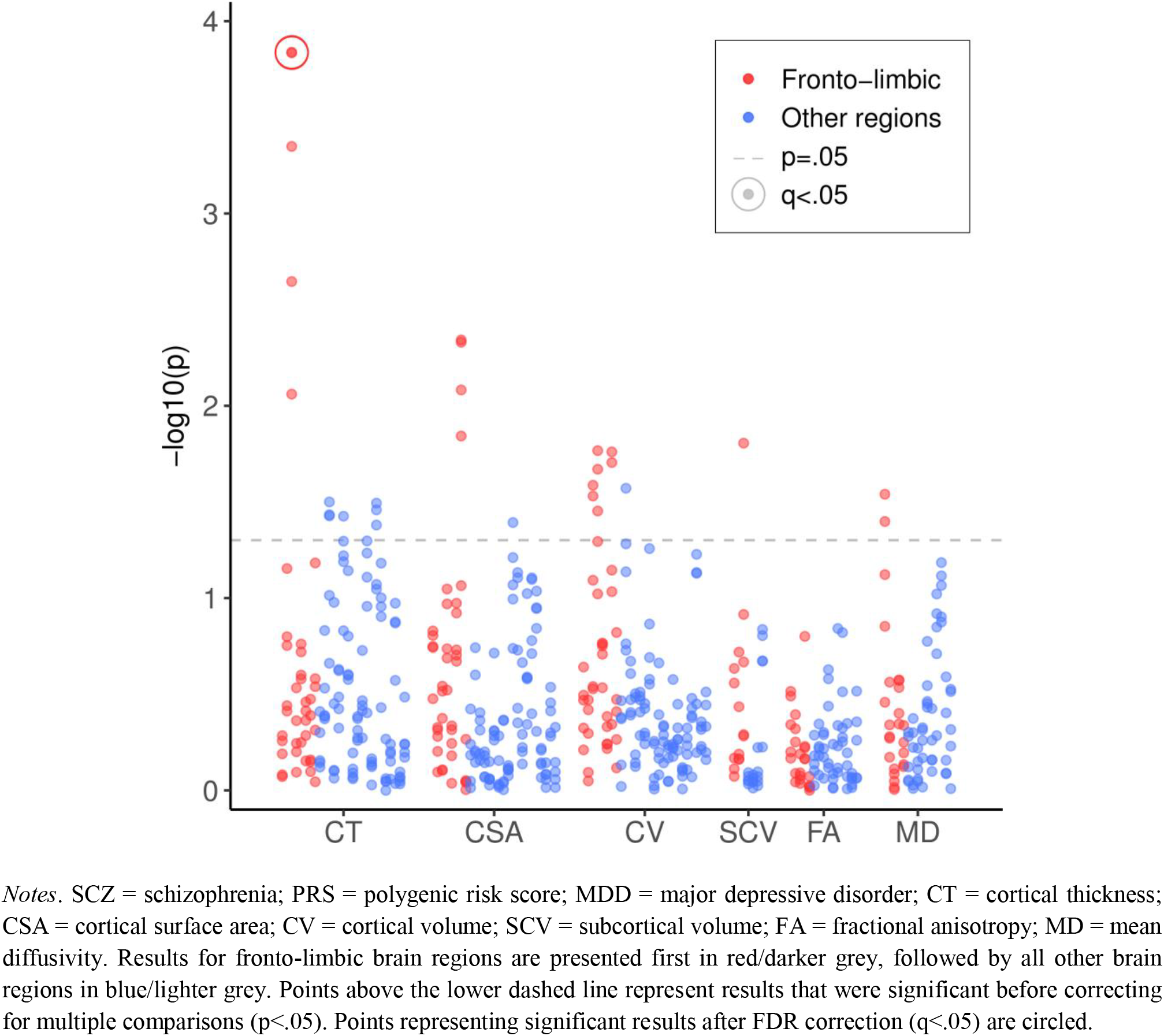
Significance of SCZ PRS by MDD interactions on all cortical, subcortical and white matter metrics

### Cortical regions

Analyses of cortical metrics focused on a subset of 1,877 subjects for whom these data were available. We first assessed interactive effects between SCZ PRS and MDD on regional mean cortical thickness. There were significant interactive effects across multiple PRS thresholds of at least β=.132 (p=.009) in rostral ACC, β=.101 (p=.042) in superior temporal gyrus and β=.120 (p=.037) in entorhinal cortex (Table 2). The interaction in rostral ACC was significant across all thresholds and the two strongest of these effects remained significant after FDR correction: for PRS threshold .50 (β=.191, q=.043) and for all SNPs (β=.191, q=.043). These interactions were driven by significant negative effects of β=–.094 (p=.001) and β=–.090 (p=.002) among controls, but positive effects of β=.103 (p=.017) and β=.109 (p=.013) among cases. These results mean that greater SCZ PRS was associated with thinner rostral ACC in controls, but not in MDD cases (Figure 2). Paired t-tests confirmed that the case-control difference was significant at PRS thresholds .05 (t_111_=2.590, p=.011) and .50 (t_111_=2.289, p=.024) among those of below-average SCZ PRS, but not at any threshold for those of above-average SCZ PRS.

**Table 2.** Interactive effects between SCZ PRS and MDD status on mean cortical thickness by region.

|  | PRS threshold .01 |  |  | PRS threshold .05 |  |  | PRS threshold .10 |  |  | PRS threshold .50 |  |  | PRS all SNPs |  |  |
| --- | --- | --- | --- | --- | --- | --- | --- | --- | --- | --- | --- | --- | --- | --- | --- |
| | $\beta$ | SE | p | $\beta$ | SE | p | $\beta$ | SE | p | $\beta$ | SE | p | $\beta$ | SE | p |
| Dorsolateral PFC | .052 | .046 | .263 | .027 | .046 | .563 | .054 | .047 | .251 | .063 | .046 | .173 | .061 | .046 | .190 |
| Caudal MFG | .043 | .046 | .347 | .018 | .046 | .703 | .041 | .047 | .386 | .036 | .046 | .433 | .030 | .046 | .518 |
| IFG | .039 | .047 | .408 | .012 | .047 | .798 | .046 | .047 | .335 | .017 | .047 | .710 | .018 | .047 | .694 |
| Lateral OFC | .032 | .049 | .519 | .023 | .050 | .646 | .030 | .050 | .553 | .010 | .050 | .848 | .011 | .050 | .830 |
| Medial OFC | .093 | .051 | .070 | .047 | .052 | .363 | .045 | .052 | .387 | .073 | .052 | .159 | .070 | .052 | .176 |
| Frontal pole | .091 | .049 | .066 | .006 | .050 | .902 | .035 | .050 | .486 | .055 | .049 | .262 | .053 | .050 | .289 |
| Rostral ACC | .132 | .050 | .009 ** | .154 | .050 | .002 ** | .179 | .051 | <.001 *** | **.191** | **.050** | **<.001 \*\*\*** | **.191** | **.050** | **<.001 \*\*\*** |
| Caudal ACC | .050 | .048 | .293 | .038 | .048 | .434 | .028 | .049 | .570 | .023 | .048 | .630 | .012 | .048 | .805 |
| Posterior cingulate | -.032 | .048 | .501 | -.042 | .048 | .389 | -.019 | .049 | .700 | -.017 | .048 | .718 | -.015 | .048 | .756 |
| Isthmus cingulate | -.040 | .048 | .408 | -.070 | .048 | .147 | -.040 | .049 | .413 | -.050 | .048 | .294 | -.038 | .048 | .424 |
| Precentral | .056 | .047 | .238 | .034 | .048 | .474 | .057 | .048 | .235 | .042 | .047 | .377 | .047 | .047 | .327 |
| Postcentral | .094 | .048 | .051 | .070 | .048 | .148 | .091 | .049 | .065 | .090 | .048 | .060 | .100 | .048 | .038 * |
| Paracentral | .087 | .048 | .072 | .054 | .049 | .265 | .069 | .049 | .158 | .055 | .048 | .253 | .056 | .049 | .249 |
| Precuneus | .022 | .046 | .639 | .008 | .046 | .870 | .013 | .047 | .774 | .011 | .046 | .812 | .009 | .046 | .852 |
| Superior parietal | .038 | .047 | .424 | .033 | .047 | .487 | .039 | .048 | .413 | .028 | .047 | .550 | .030 | .047 | .533 |
| Inferior parietal | .039 | .045 | .385 | .028 | .045 | .533 | .060 | .045 | .187 | .043 | .045 | .341 | .041 | .045 | .362 |
| Supramarginal | .074 | .046 | .110 | .040 | .047 | .395 | .083 | .047 | .078 | .088 | .046 | .058 | .091 | .046 | .050 |
| Insula | .004 | .051 | .937 | -.016 | .051 | .753 | .019 | .052 | .711 | -.021 | .051 | .685 | -.017 | .051 | .737 |
| STG | .101 | .050 | .042 * | .108 | .050 | .032 * | .107 | .051 | .035 * | .085 | .050 | .090 | .087 | .050 | .085 |
| MTG | .043 | .049 | .371 | .081 | .049 | .100 | .091 | .050 | .066 | .075 | .049 | .125 | .078 | .049 | .111 |
| ITG | .007 | .053 | .896 | .009 | .053 | .869 | .033 | .054 | .541 | .007 | .053 | .899 | .000 | .053 | >.999 |
| Fusiform | -.020 | .054 | .706 | -.030 | .054 | .575 | -.009 | .055 | .874 | -.027 | .055 | .628 | -.025 | .055 | .645 |
| Entorhinal | .070 | .057 | .218 | .096 | .058 | .097 | .125 | .058 | .032 * | .120 | .057 | .037 * | .120 | .057 | .037 * |
| Parahippocampal | -.083 | .051 | .105 | -.048 | .052 | .354 | -.009 | .052 | .864 | .013 | .052 | .794 | .014 | .052 | .785 |
| Temporal pole | -.005 | .050 | .922 | .056 | .050 | .268 | .076 | .051 | .135 | .081 | .050 | .106 | .075 | .050 | .133 |
| Lateral occipital | -.004 | .046 | .925 | -.006 | .047 | .895 | .012 | .047 | .802 | -.008 | .047 | .861 | -.009 | .047 | .850 |
| Medial occipital | -.027 | .048 | .577 | -.048 | .049 | .327 | -.028 | .049 | .573 | -.023 | .048 | .634 | -.020 | .049 | .673 |

*Notes.* SCZ = schizophrenia; PRS = polygenic risk score; MDD = major depressive disorder; PFC = prefrontal cortex; MFG = middle frontal gyrus; IFG = inferior frontal gyrus; OFC = orbitofrontal cortex; ACC = anterior cingulate cortex; STG = superior temporal gyrus; MTG = middle temporal gyrus; ITG = inferior temporal gyrus. Reported results are for interactive effects between SCZ PRS and MDD status on regional mean cortical thickness, controlling covariates. \*, \*\* and \*\*\* represent significant results at $p < .05$ , $p < .01$ and $p < .001$ , respectively, before correcting for multiple comparisons. Significant results after FDR correction are highlighted in bold.

**Figure 2.**
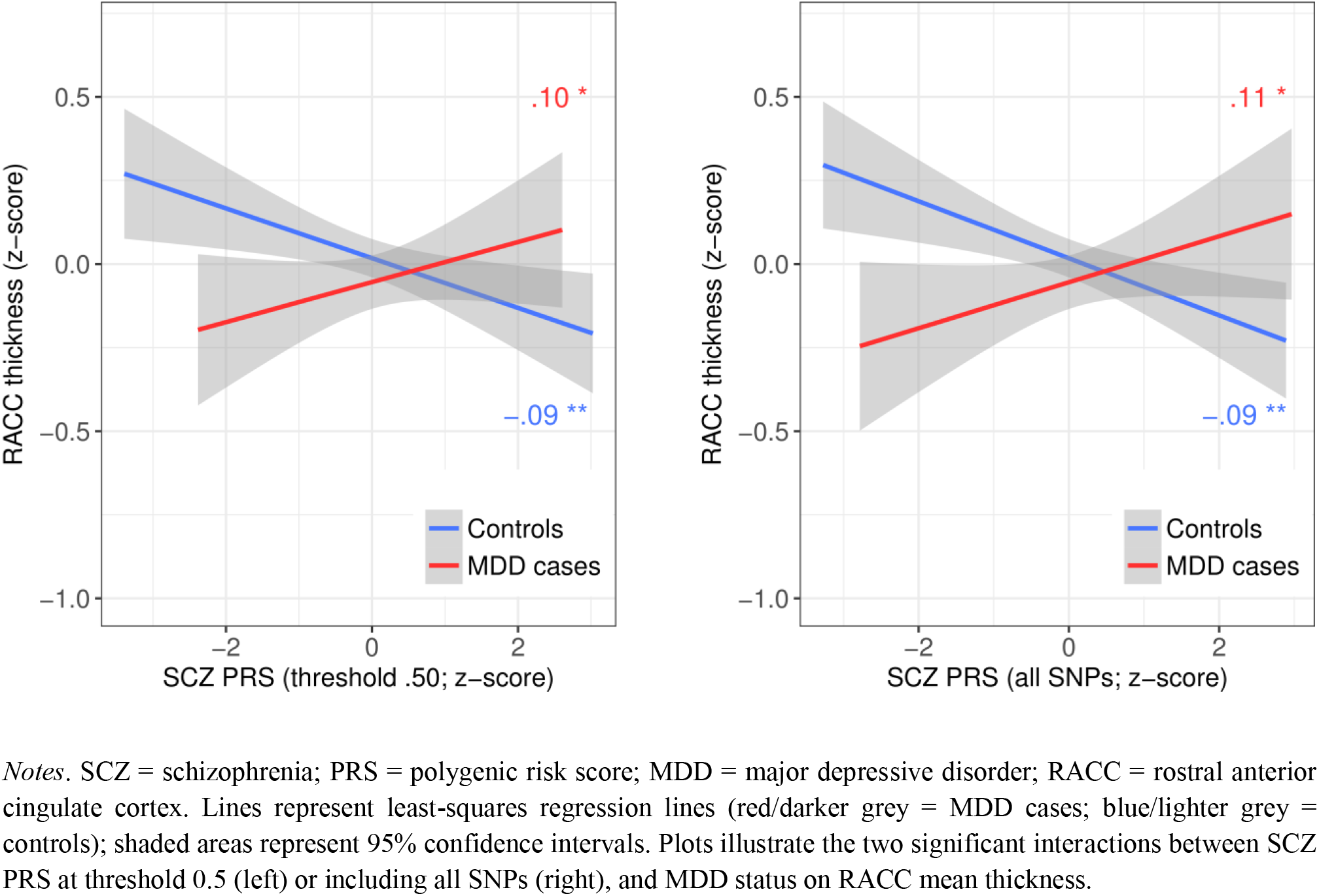
Mean thickness by SCZ PRS and MDD group for the two significant SCZ PRS by MDD interactions in rostral anterior cingulate cortex

Given the specific effect of SCZ PRS on rostral ACC mean thickness among MDD cases, we tested whether it was influenced by a subgroup of participants. Repeated k-means clustering of SCZ PRS and rostral ACC thickness data using the function “pamk” identified three as the optimum (according to average silhouette width; also verified using hierarchical clustering) for each of the three thresholds showing significant interactions. However, as shown in supplementary *Figure S1*, for each threshold, none of the three clusters clearly represented a high-SCZ-risk subgroup, and none was significantly distinct from the other two, with low Dunn indices (DIs) of .009.

As more cortical data became available later, we were able to retest the interaction between SCZ PRS and MDD on rostral ACC thickness in a replication sample. The interaction was weaker but significant for SCZ PRS thresholds .05 (β=.095, p=.024) and .10 (β=.083, p=.047). As shown in supplementary *Figure S2*, the interaction was attenuated mainly by the controls of the replication sample, among whom the negative association between SCZ PRS and rostral ACC thickness was much closer to zero than in the discovery sample (β=–.013, p=.595; β=–.009, p=.724). The association among MDD cases of the replication sample was still positive, but also weaker than in the discovery sample and no longer significant (β=.065, p=.074; β=.057, p=.116).

For cortical surface area, there were significant interactions across multiple PRS thresholds of β=–.091 (p=.014) to β=–.105 (p=.005) in inferior frontal gyrus (IFG), again due to a stronger negative effect of SCZ PRS among MDD cases than controls, but these results failed to survive FDR correction. Full results are reported in *Table S1*. For cortical volume, there were significant interactions across multiple PRS thresholds of β=–.088 (p=.020) and β=–.090 (p=.017) for IFG, β=.091 (p=.029) and β=.094 (p=.026) for rostral ACC; and β=.098 (p=.035) to β=.110 (p=.017) for caudal ACC. However, following FDR correction, none of these interactive effects on regional cortical volumes remained significant (*Table S2*).

### Subcortical structures

We next tested interactive effects on subcortical volumes, which were available for all 4,944 subjects. Across all subcortical volumes and PRS thresholds, there was only one significant SCZ PRS by MDD interaction: a weak positive interactive effect of β=.047 (p=.016) on thalamic volume at PRS threshold .01 (*Table S3*). This was driven by weak and non-significant but opposite associations between SCZ PRS and thalamic volume in MDD cases (β=.030) and controls (β=–.022), but the interaction did not remain significant after FDR correction.

### White matter tracts

Finally, we explored interactions in white matter tracts for the 4,420 subjects with DTI data. There were no significant interactive effects between SCZ PRS and MDD on the FA of any of the white matter tracts at any of the PRS thresholds (*Table S4*). For MD, there was a significant interactive effect of β=.057 (p=.045) at threshold .50 and β=.061 (p=.033) with all SNPs in the uncinate fasciculus (*Table S5*). Uncinate MD also showed weak and non-significant opposite effects of SCZ PRS in MDD cases (β=.037; β=.035) and controls (β=–.024; β=–.030). No results survived correction for multiple comparisons.

## Discussion

Following on from findings that MDD differs behaviourally as a function of genetic risk for SCZ, we tested interactions between SCZ PRS and MDD status on a range of brain structure measures in a large sample of UK Biobank participants. Interactions were not significant for most brain measures, but results for rostral ACC mean thickness at two PRS thresholds remained significant after correction for multiple comparisons. In rostral ACC, greater SCZ PRS showed a weak association with thinner cortex among controls, but a weak association with *thicker* cortex among MDD cases. However, MDD cases and controls only completely diverged toward the lower end of the range in SCZ PRS, where cases still showed thinner rostral ACC. We were able to replicate this interaction in a second subsample, although the effect was attenuated. This significant result seems to indicate localised reserve or protection against the negative neurostructural effects of SCZ-associated genes among individuals with MDD, but alternative explanations are discussed below.

We focused on interactions between SCZ PRS and MDD, but also tested main effects of SCZ PRS within MDD case and control groups for significant interactions. For controls, results suggested that a higher SCZ PRS was associated with reductions in rostral ACC thickness, as expected. These results are consistent with the reductions in prefrontal volumes seen in SCZ (Ross et al., 2006; Keshavan et al., 2008), and therefore with previous evidence of schizophrenia-like neuromorphology among high-risk but healthy controls (Lawrie et al., 2001; Boos et al., 2007). Further, case-control t-tests clearly showed deficits in rostral ACC thickness among MDD cases with a low SCZ PRS, compared to low-SCZ-risk controls. This is also expected and consistent with previous evidence of reduced cingulate grey matter (Salvadore et al., 2011; Rodríguez-Cano et al., 2014; Schmaal et al., 2017) in MDD. However, others have observed increased thickness of rostral ACC in MDD (Ancelin et al., 2019) and of other cortical regions in SCZ (Dukart et al., 2017), which does correspond to our findings among individuals at higher genetic risk of SCZ.

Increased rostral ACC thickness among individuals with both MDD and a high genetic predisposition to SCZ may initially seem counterintuitive, suggesting that two putative neuropathologies instead confer benefits when combined. However, while positive associations between SCZ PRS and brain measures among individuals with MDD also stand in contrast with most previous findings (as above), some previous findings are comparable. For example, Papmeyer et al. (2015) reported thickening of inferior frontal and precentral cortices among individuals with both MDD and familial risk of bipolar disorder. One possible explanation is that the subset of MDD cases with a higher SCZ PRS in this study included individuals who had not developed SCZ *despite* a high level of genetic predisposition. These individuals may have had a neurobiological advantage over others at high genetic risk who *did* develop SCZ, who were not included in the study. Resistance to aberrant neurodevelopmental influences could explain both why high-SCZ-risk MDD cases did not develop SCZ and why they were more comparable to controls than to low-SCZ-risk cases. Or, simply, the greater rostral ACC thickness observed among high-SCZ-risk MDD cases may have protected against more severe psychopathology. However, as only a small proportion of individuals develop SCZ – even among those with a higher PRS – their exclusion is unlikely to have had such an impact on our results.

Alternatively, it may be that the increase in rostral ACC thickness that we observed does not actually represent a benefit. Increased thickness and functional connectivity of rostral ACC have been associated with insomnia (Winkelman et al., 2013) and delusions in SCZ (Schott et al., 2015), respectively. Such symptoms may relate to a deficit in, for example, normal axonal pruning, as this could lead to increased cortical thickness. Some theories of SCZ associate the disorder with altered pruning throughout development (Feinberg, 1982; Keshavan et al., 1994; Glausier & Lewis; 2013), although they typically attribute it to excessive pruning, rather than decreased pruning. However, disruption of normal neurodevelopmental mechanisms could lead to excessive pruning in some areas but decreased pruning in others. While this could in turn lead to greater cortical thickness, these increases might be associated with aberrant hyperactivity or reduced efficiency, contributing to psychiatric symptoms. Further results do suggest SCZ-related increases in thickness and volume of specific brain regions, such as superior parietal cortex and the amygdala (Spoletini et al.; 2011; van Haren et al., 2011), supporting this account. Increases in ACC thickness have also been observed with ageing in some studies (Salat et al., 2004; Abe et al., 2008). Whether our results relate to a benefit or impairment specifically among high-SCZ-risk MDD cases, they provide some evidence of an SCZ-risk-related MDD subtype distinct in terms of prefrontal morphology.

Considering that MDD cases and controls only completely diverged at the lower end of the SCZ PRS scale, it may be that case-control differences in rostral ACC thickness were not reversed at the higher end of the range in SCZ PRS, but merely attenuated. In this case, the results relate more directly to Whalley et al.’s (2016) findings of attenuated case-control differences in neuroticism and psychological distress at higher SCZ PRS. This rekindles the idea that both sets of findings represent a protective effect of genetic risk for SCZ among MDD cases, and although it remains unclear why SCZ risk might be protective, the correspondence between clinical and neuroimaging findings strengthens evidence for a possible SCZ-risk-related MDD subtype. Although our cluster analyses were not able to distinguish this subtype, it warrants further investigation.

For most other structural brain metrics assessed, however, we found no significant influence of SCZ PRS on MDD case-control differences. This may have been due to limitations of the study. We used data from UK Biobank, which recruited large numbers of participants from the general population, but our sample (and particularly the smaller subset with data on cortical metrics) may not have been large enough to detect subtle genetic effects. The generalizability of results from UK Biobank is also still potentially limited by selection bias (Allen et al., 2012), as the sample is healthier overall than the general population (Fry et al., 2017). This is a common limitation to most observational studies but, as Fry et al. argue, results are likely to be both robust and widely generalizable. However, a limitation of recruiting from the general population is that SCZ PRS may not have been particularly high for any of our subjects. We still found significant heterogeneity in MDD related to SCZ PRS despite this, which perhaps lends further weight to the conclusion drawn from our significant results for rostral ACC thickness. Investigation of similar effects in a sample including a group of individuals with a higher PRS for SCZ (e.g. relatives of patients) may reveal more about the potential SCZ-risk-related MDD subtype.

UK Biobank participants are also older than the general population, with an average age of over 60 years, although a range of age groups from middle to late adulthood are represented. Focusing on this age group did, however, make it unlikely that subjects were yet to be diagnosed with SCZ, meaning that our results do relate to the effects of genetic risk only, rather than to SCZ itself. Another possible limitation was our measure of MDD, as it was derived from a web-based questionnaire. However, the online CIDI-SF was the most reliable method that could be efficiently used in such a large sample, and provided clinically relevant information based on DSM-IV diagnostic criteria. Finally, there were a number of potential covariates that we were unable to include, as complete data were unavailable at the time of analysis, for example, subjects’ use of alcohol, tobacco, other drugs and medications, age of disorder onset and disorder severity. This should be considered when interpreting the current findings.

In summary, we tested for the presence of a distinct subtype of MDD associated with high genetic risk for SCZ by exploring interactive effects between these two variables on structural measures across the brain. Results were generally not significant, but we did observe a significant interactive effect on cortical thickness in rostral ACC for two PRS thresholds, which we were also able to replicate in a second subsample. While the direction of this interaction was unexpected, our results indicate that differences between MDD cases and controls in rostral ACC thickness are significantly influenced by SCZ PRS. This finding, together with related previous findings, suggest subtle differences in MDD related to SCZ risk and rostral ACC, which could influence important factors such as development of the disorder and response to treatment.

## Acknowledgements

This study was supported by a Wellcome Trust Strategic Award, “Stratifying Resilience and Depression Longitudinally” (STRADL; ref. 104036/Z/14/Z) and conducted using the UK Biobank Resource (application no. 4844). We would like to thank UK Biobank participants and researchers for their contributions to the study. Part of the work was undertaken at The Centre for Cognitive Ageing and Cognitive Epidemiology (CCACE), part of the cross-council Lifelong Health and Wellbeing Initiative (MR/K026992/1); funding from the Medical Research Council (MRC) and Biotechnology and Biological Sciences Research Council (BBSRC) is gratefully acknowledged. Age UK also provided support for the work undertaken at CCACE (via the Disconnected Mind project). We would also like to acknowledge David Liewald of CCACE for his part in processing imaging data. MAH is supported by The Dr Mortimer and Theresa Sackler Foundation. SRC is supported by the MRC (grants MR/M013111/1 and MR/R024065/1) and the National Institutes of Health (R01AG054628). HCW is supported by a John, Margaret, Alfred and Stewart (JMAS) Sim fellowship from the Royal College of Physicians of Edinburgh (RCPE), and an Edinburgh Scientific Academic Track (ESAT) fellowship from the University of Edinburgh.

