## Supplementary material for "Stratifying major depressive disorder by polygenic risk for schizophrenia in relation to structural brain measures"

### Supplementary Methods

#### Brain imaging data

*Acquisition:* All brain magnetic resonance imaging (MRI) data were acquired at the same site, using the same Siemens (Berlin/Munich, Germany) Skyra 3T scanner and 32-channel head coil, and the same protocol (Miller et al., 2016). Briefly, T1 structural data were acquired using a 3D magnetization-prepared rapid acquisition gradient-echo (MPRAGE) sequence, at 1mm isotropic resolution, and diffusion imaging data were acquired using a standard monopolar Stejskal-Tanner sequence, at 2mm spatial resolution, with two b-values of 1000 and 2000 s/mm$^2$. Raw imaging data were processed and quality-assessed using an automated pipeline by the UK Biobank imaging team (Alfaro-Almagro et al., 2018) and resulting imaging-derived phenotypes were made available to approved researchers.

*Cortical regions:* Cortical measures were derived locally using FreeSurfer version 5.3 (Dale et al., 1999; Fischl et al., 1999; Fischl et al., 2004) to segment the cortex according to the Desikan-Killany atlas (Desikan et al., 2006) into 34 regions per hemisphere. Output data were visually assessed, excluding data for subjects with major errors (~10%) and for regions with localised errors (~2%). Metrics for 11 regions per hemisphere were weighted-averaged (mean thickness) or summed (surface area and volume) to produce metrics for four larger and more commonly studied regions – dorsolateral prefrontal cortex, inferior frontal gyrus, superior temporal gyrus, and medial occipital cortex – as in previous studies (Cox et al., 2018). After ruling out hemispheric effects, left and right metrics for each region were also combined, resulting in 27 bilateral regions. At the time of the present study, FreeSurfer processing and quality control had been completed for only a subset of the available neuroimaging sample (N=2,966).

*Subcortical structures:* T1 images were processed by the UK Biobank imaging team, which included the segmentation of the left and right caudate nucleus, thalamus, putamen, pallidum, hippocampus, amygdala and nucleus accumbens using FIRST (Patenaude et al., 2011). Volumes of subcortical structures were available for all 7,536 subjects and were summed bilaterally prior to further analysis.

*White matter tracts:* The diffusion tensor imaging (DTI) model, fitted voxel-wise during acquisition of diffusion images, provided measures of fractional anisotropy (FA) and mean diffusivity (MD). As per the automated processing pipeline (Alfaro-Almagro et al., 2018), within-voxel modelling and probabilistic tractography were performed using bedpostx and probtrackx (Behrens et al., 2007), generating measures for 27 specific tracts defined using AutoPtx (de Groot et al., 2013), 24 of which were combined bilaterally prior to further analysis. These tract-averaged measures of white matter microstructural integrity were available for 6,677 subjects.

**Figure S1 – Clusters derived from SCZ PRS and rostral anterior cingulate cortex mean thickness for the two significant SCZ PRS by MDD interactions.**

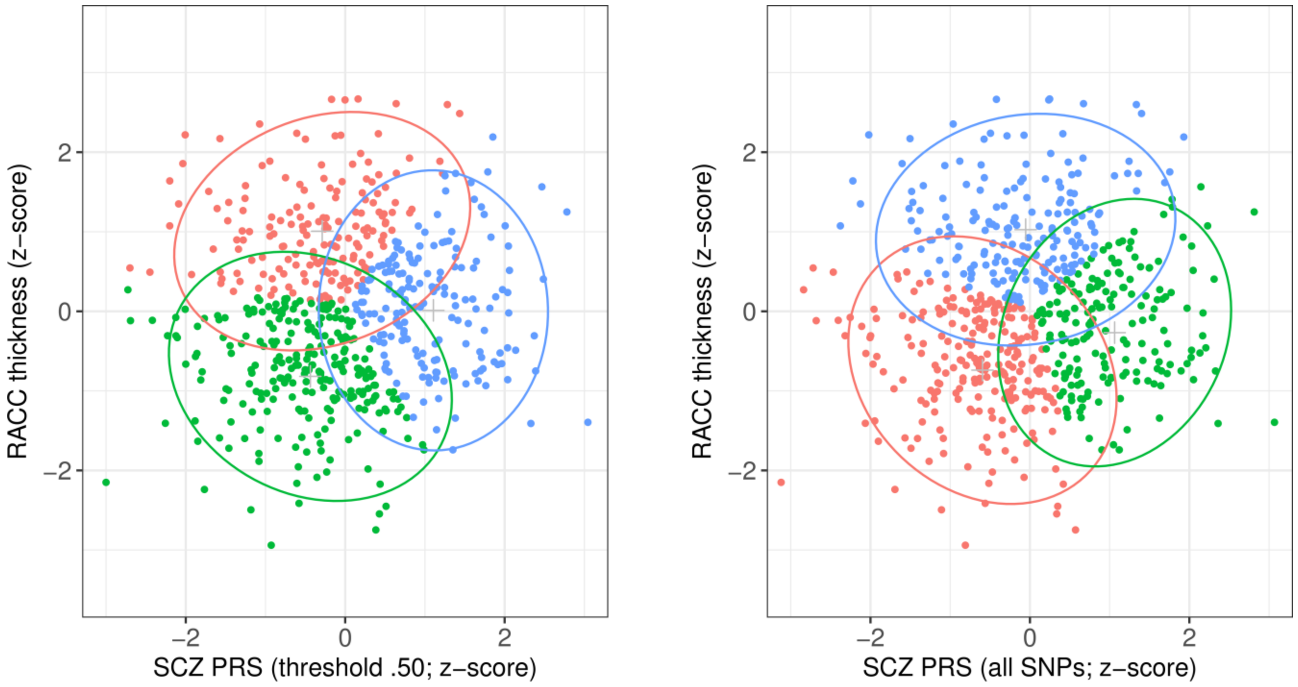


*Notes*. SCZ = schizophrenia; PRS = polygenic risk score; MDD = major depressive disorder; RACC = rostral anterior cingulate cortex. Red/green/blue points represents subjects included in each of the three derived clusters; surrounding ellipses represent 95% confidence intervals. Plots illustrate clusters derived from data relating to the two significant interactions between SCZ PRS at threshold 0.5 (left) or including all SNPs (right), and MDD status on RACC mean thickness.

**Figure S2 – Mean rostral anterior cingulate cortex thickness by SCZ PRS and MDD group for the two significant SCZ PRS by MDD interactions in the replication sample.**

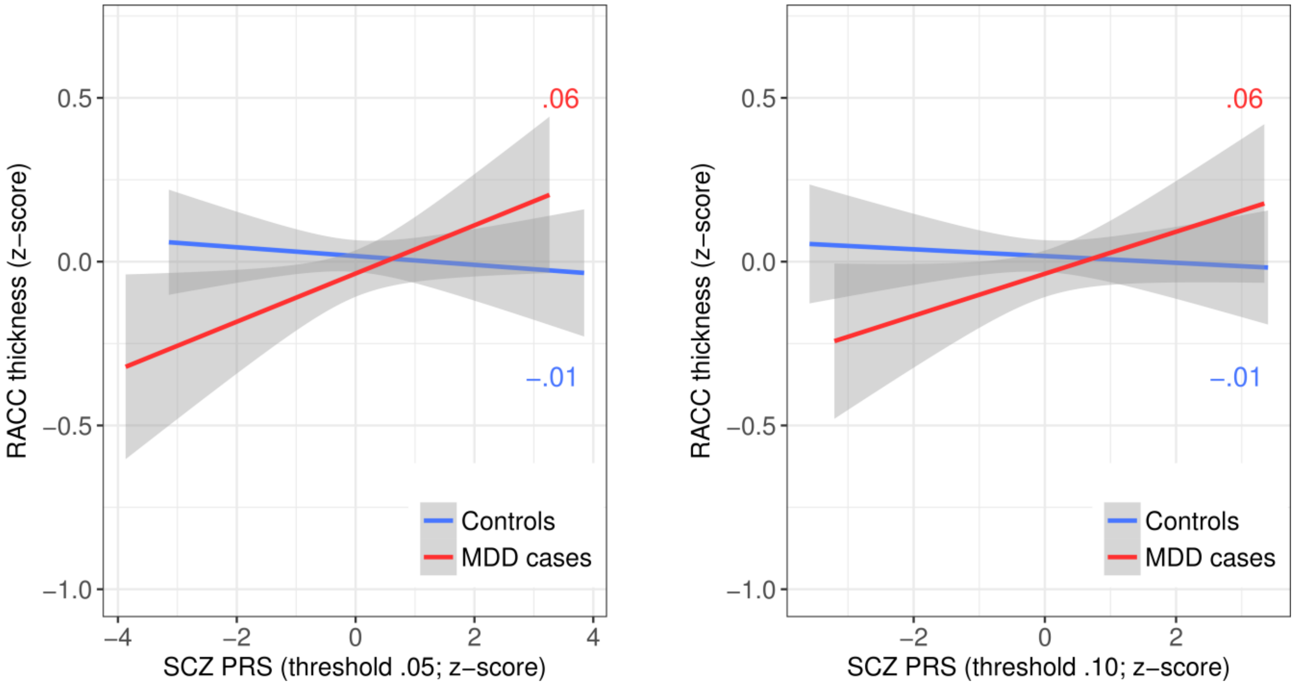


*Notes*. SCZ = schizophrenia; PRS = polygenic risk score; MDD = major depressive disorder; RACC = rostral anterior cingulate cortex. Lines represent least-squares regression lines (red/darker grey = MDD cases; blue/lighter grey = controls); shaded areas represent 95% confidence intervals. Plots illustrate the two significant interactions between SCZ PRS at threshold 0.5 (left) or including all SNPs (right), and MDD status on RACC mean thickness in the replication sample.

**Table S1 – Interactive effects between SCZ PRS and MDD status on cortical surface area by region.**

| | PRS threshold .01 | | | PRS threshold .05 | | | PRS threshold .10 | | | PRS threshold .50 | | | PRS all SNPs | | |
|---|---|---|---|---|---|---|---|---|---|---|---|---|---|---|---|
| | β | SE | p | β | SE | p | β | SE | p | β | SE | p | β | SE | p |
| Dorsolateral PFC | -.003 | .026 | .918 | .011 | .026 | .661 | .015 | .026 | .570 | .018 | .026 | .484 | .019 | .026 | .462 |
| Caudal MFG | -.061 | .038 | .106 | -.050 | .038 | .186 | -.060 | .038 | .120 | -.049 | .038 | .198 | -.047 | .038 | .213 |
| IFG | -.063 | .037 | .086 | -.091 | .037 | .014 * | -.099 | .038 | .008 ** | -.105 | .037 | .005 ** | -.105 | .037 | .005 ** |
| Lateral OFC | -.045 | .033 | .178 | -.048 | .033 | .148 | -.033 | .034 | .334 | -.045 | .033 | .181 | -.047 | .033 | .157 |
| Medial OFC | -.024 | .034 | .471 | -.009 | .034 | .803 | .017 | .035 | .627 | .024 | .034 | .485 | .022 | .034 | .524 |
| Frontal pole | -.027 | .045 | .542 | -.006 | .045 | .889 | -.005 | .046 | .909 | -.006 | .045 | .900 | -.001 | .045 | .991 |
| Rostral ACC | .010 | .039 | .788 | .011 | .039 | .781 | .032 | .039 | .422 | .040 | .039 | .305 | .041 | .039 | .287 |
| Caudal ACC | .057 | .043 | .183 | .044 | .043 | .301 | .055 | .043 | .205 | .073 | .043 | .090 | .069 | .043 | .107 |
| Posterior cingulate | .033 | .037 | .377 | -.001 | .037 | .970 | .005 | .038 | .896 | .021 | .037 | .575 | .017 | .037 | .655 |
| Isthmus cingulate | -.013 | .036 | .716 | -.018 | .036 | .623 | -.017 | .036 | .636 | -.048 | .036 | .181 | -.041 | .036 | .251 |
| Precentral | -.012 | .032 | .709 | -.005 | .032 | .870 | -.005 | .032 | .876 | .005 | .032 | .875 | -.002 | .032 | .938 |
| Postcentral | -.044 | .034 | .193 | -.024 | .034 | .488 | -.022 | .034 | .520 | -.023 | .034 | .492 | -.020 | .034 | .548 |
| Paracentral | .025 | .040 | .521 | .015 | .040 | .700 | -.002 | .040 | .964 | .003 | .040 | .944 | -.003 | .040 | .931 |
| Precuneus | -.026 | .033 | .430 | -.026 | .033 | .434 | .000 | .033 | .991 | -.004 | .033 | .915 | -.004 | .033 | .899 |
| Superior parietal | -.010 | .036 | .783 | .007 | .037 | .840 | .012 | .037 | .742 | -.011 | .036 | .766 | -.010 | .037 | .783 |
| Inferior parietal | -.047 | .035 | .183 | -.058 | .035 | .101 | -.073 | .036 | .040 * | -.066 | .035 | .062 | -.061 | .035 | .085 |
| Supramarginal | -.027 | .035 | .445 | -.031 | .035 | .374 | -.047 | .036 | .186 | -.062 | .035 | .078 | -.063 | .035 | .074 |
| Insula | .042 | .034 | .217 | .023 | .034 | .511 | .012 | .035 | .729 | .018 | .034 | .599 | .017 | .034 | .628 |
| STG | -.023 | .031 | .454 | -.035 | .031 | .260 | -.053 | .032 | .095 | -.035 | .031 | .263 | -.035 | .031 | .257 |
| MTG | -.030 | .035 | .399 | -.063 | .036 | .080 | -.064 | .036 | .079 | -.049 | .036 | .166 | -.046 | .036 | .194 |
| ITG | -.027 | .039 | .493 | -.062 | .039 | .112 | -.058 | .040 | .144 | -.067 | .040 | .092 | -.063 | .040 | .113 |
| Fusiform | .020 | .040 | .605 | -.007 | .040 | .858 | -.020 | .040 | .619 | -.014 | .040 | .732 | -.014 | .040 | .719 |
| Entorhinal | .038 | .049 | .432 | .034 | .049 | .486 | .042 | .050 | .395 | .019 | .049 | .697 | .023 | .049 | .645 |
| Parahippocampal | .015 | .041 | .718 | -.008 | .042 | .848 | -.008 | .042 | .856 | -.020 | .042 | .630 | -.018 | .042 | .661 |
| Temporal pole | .010 | .044 | .826 | -.007 | .045 | .879 | .016 | .045 | .718 | .002 | .045 | .965 | .010 | .045 | .825 |
| Lateral occipital | -.031 | .036 | .387 | -.023 | .036 | .523 | -.038 | .036 | .291 | -.033 | .036 | .350 | -.024 | .036 | .502 |
| Medial occipital | -.028 | .038 | .470 | -.002 | .038 | .965 | -.014 | .039 | .715 | .006 | .038 | .877 | .010 | .038 | .800 |

*Notes*. SCZ = schizophrenia; PRS = polygenic risk score; MDD = major depressive disorder; PFC = prefrontal cortex; MFG = middle frontal gyrus; IFG = inferior frontal gyrus; OFC = orbitofrontal cortex; ACC = anterior cingulate cortex; STG = superior temporal gyrus; MTG = middle temporal gyrus; ITG = inferior temporal gyrus. Reported results are for interactive effects between SCZ PRS and MDD status on regional cortical surface area, controlling covariates. * and ** represent significant results at $p<.05$ and $p<.01$, respectively, before correcting for multiple comparisons. No results remained significant after FDR correction.

**Table S2 – Interactive effects between SCZ PRS and MDD status on cortical volume by region.**

| | PRS threshold .01 | | | PRS threshold .05 | | | PRS threshold .10 | | | PRS threshold .50 | | | PRS all SNPs | | |
|---|---|---|---|---|---|---|---|---|---|---|---|---|---|---|---|
| | β | SE | p | β | SE | p | β | SE | p | β | SE | p | β | SE | p |
| Dorsolateral PFC | .036 | .027 | .196 | .029 | .028 | .293 | .038 | .028 | .177 | .038 | .028 | .173 | .038 | .028 | .171 |
| Caudal MFG | -.028 | .039 | .467 | -.020 | .039 | .606 | -.032 | .040 | .413 | -.022 | .039 | .573 | -.022 | .039 | .576 |
| IFG | -.028 | .038 | .453 | -.068 | .038 | .072 | -.064 | .038 | .093 | -.088 | .038 | .020 * | -.090 | .038 | .017 * |
| Lateral OFC | -.026 | .036 | .474 | -.043 | .036 | .228 | -.018 | .036 | .616 | -.034 | .036 | .339 | -.036 | .036 | .320 |
| Medial OFC | .009 | .038 | .806 | .005 | .038 | .893 | .026 | .038 | .506 | .036 | .038 | .338 | .033 | .038 | .381 |
| Frontal pole | .069 | .048 | .151 | .015 | .049 | .764 | .030 | .049 | .543 | .042 | .048 | .390 | .047 | .048 | .336 |
| Rostral ACC | .044 | .042 | .297 | .045 | .042 | .288 | .074 | .043 | .081 | .091 | .042 | .029 * | .094 | .042 | .026 * |
| Caudal ACC | .110 | .046 | .017 * | .078 | .047 | .095 | .092 | .047 | .051 | .107 | .046 | .021 * | .098 | .046 | .035 * |
| Posterior cingulate | .038 | .040 | .342 | -.002 | .040 | .954 | .014 | .040 | .737 | .032 | .040 | .417 | .032 | .040 | .425 |
| Isthmus cingulate | -.050 | .038 | .187 | -.074 | .038 | .052 | -.052 | .038 | .173 | -.084 | .038 | .027 * | -.068 | .038 | .073 |
| Precentral | .031 | .036 | .392 | .024 | .037 | .518 | .037 | .037 | .324 | .037 | .036 | .308 | .035 | .036 | .338 |
| Postcentral | .025 | .039 | .523 | .022 | .039 | .573 | .037 | .039 | .353 | .026 | .039 | .503 | .034 | .039 | .375 |
| Paracentral | .079 | .041 | .055 | .062 | .041 | .137 | .053 | .042 | .204 | .046 | .041 | .264 | .045 | .041 | .280 |
| Precuneus | -.019 | .033 | .555 | -.019 | .033 | .572 | .006 | .033 | .865 | .001 | .033 | .985 | -.002 | .033 | .947 |
| Superior parietal | .008 | .037 | .830 | .031 | .038 | .404 | .047 | .038 | .218 | .015 | .037 | .682 | .018 | .038 | .632 |
| Inferior parietal | -.013 | .036 | .719 | -.021 | .036 | .558 | -.027 | .037 | .468 | -.027 | .036 | .459 | -.024 | .036 | .515 |
| Supramarginal | .015 | .036 | .679 | -.005 | .037 | .896 | -.005 | .037 | .899 | -.020 | .037 | .583 | -.019 | .037 | .603 |
| Insula | .040 | .036 | .265 | .016 | .036 | .667 | .020 | .037 | .595 | .019 | .036 | .605 | .020 | .036 | .574 |
| STG | .032 | .035 | .356 | .026 | .036 | .473 | .018 | .036 | .620 | .022 | .035 | .544 | .022 | .036 | .543 |
| MTG | -.001 | .038 | .983 | -.010 | .038 | .794 | -.008 | .038 | .845 | .011 | .038 | .779 | .012 | .038 | .760 |
| ITG | -.012 | .042 | .780 | -.030 | .043 | .474 | -.020 | .043 | .650 | -.027 | .043 | .535 | -.023 | .043 | .585 |
| Fusiform | -.008 | .043 | .851 | -.042 | .043 | .332 | -.039 | .044 | .374 | -.035 | .043 | .419 | -.036 | .043 | .409 |
| Entorhinal | .042 | .051 | .406 | .060 | .052 | .247 | .065 | .052 | .213 | .044 | .052 | .393 | .050 | .052 | .337 |
| Parahippocampal | -.046 | .046 | .314 | -.045 | .046 | .328 | -.015 | .047 | .751 | -.021 | .046 | .653 | -.017 | .046 | .719 |
| Temporal pole | .023 | .049 | .636 | .046 | .050 | .355 | .090 | .050 | .074 | .088 | .049 | .074 | .093 | .049 | .059 |
| Lateral occipital | -.027 | .037 | .466 | -.026 | .037 | .482 | -.019 | .037 | .613 | -.028 | .037 | .440 | -.020 | .037 | .591 |
| Medial occipital | -.040 | .039 | .308 | -.029 | .039 | .465 | -.037 | .040 | .360 | -.019 | .039 | .635 | -.016 | .039 | .692 |

*Notes*. SCZ = schizophrenia; PRS = polygenic risk score; MDD = major depressive disorder; PFC = prefrontal cortex; MFG = middle frontal gyrus; IFG = inferior frontal gyrus; OFC = orbitofrontal cortex; ACC = anterior cingulate cortex; STG = superior temporal gyrus; MTG = middle temporal gyrus; ITG = inferior temporal gyrus. Reported results are for interactive effects between SCZ PRS and MDD status on regional cortical volume, controlling covariates. * represents significant results at $p<.05$ before correcting for multiple comparisons. No results remained significant after FDR correction.

**Table S3 – Interactive effects between SCZ PRS and MDD status on volumes of subcortical structures.**

| | PRS threshold .01 | | | PRS threshold .05 | | | PRS threshold .10 | | | PRS threshold .50 | | | PRS all SNPs | | |
|---|---|---|---|---|---|---|---|---|---|---|---|---|---|---|---|
| | β | SE | p | β | SE | p | β | SE | p | β | SE | p | β | SE | p |
| Caudate volume | .003 | .025 | .905 | -.001 | .025 | .972 | .004 | .025 | .863 | .005 | .025 | .831 | .003 | .025 | .898 |
| Thalamus volume | .047 | .019 | .016 * | .029 | .019 | .131 | .024 | .019 | .225 | .012 | .019 | .536 | .012 | .019 | .546 |
| Putamen volume | .006 | .023 | .802 | -.012 | .023 | .597 | .004 | .023 | .849 | -.002 | .023 | .943 | -.003 | .023 | .892 |
| Pallidum volume | -.014 | .026 | .584 | -.038 | .026 | .138 | -.033 | .026 | .205 | -.033 | .026 | .203 | -.037 | .026 | .149 |
| Amygdala volume | .006 | .028 | .832 | .009 | .028 | .749 | .012 | .028 | .664 | .031 | .028 | .264 | .034 | .028 | .221 |
| Accumbens volume | .004 | .026 | .892 | -.005 | .027 | .837 | -.003 | .027 | .921 | -.004 | .026 | .885 | -.007 | .026 | .800 |
| Hippocampus volume | -.009 | .027 | .731 | -.022 | .027 | .421 | -.033 | .027 | .215 | -.008 | .027 | .754 | -.010 | .027 | .716 |

*Notes*. SCZ = schizophrenia; PRS = polygenic risk score; MDD = major depressive disorder. Reported results are for interactive effects between SCZ PRS and MDD status on subcortical volumes, controlling covariates. * represents a significant result at $p<.05$ before correcting for multiple comparisons. No results remained significant after FDR correction.

**Table S4 – Interactive effects between SCZ PRS and MDD status on FA by white matter tract.**

| | PRS threshold .01 | | | PRS threshold .05 | | | PRS threshold .10 | | | PRS threshold .50 | | | PRS all SNPs | | |
|---|---|---|---|---|---|---|---|---|---|---|---|---|---|---|---|
| | β | SE | p | β | SE | p | β | SE | p | β | SE | p | β | SE | p |
| Superior longitudinal | .000 | .031 | .999 | -.006 | .031 | .846 | .002 | .031 | .958 | .002 | .031 | .945 | -.001 | .031 | .966 |
| Inferior longitudinal | -.007 | .031 | .818 | -.018 | .031 | .575 | -.005 | .031 | .874 | .001 | .031 | .965 | -.001 | .031 | .975 |
| Cingulum cingulate | .013 | .032 | .685 | -.044 | .031 | .158 | -.017 | .031 | .591 | -.013 | .031 | .685 | -.016 | .031 | .600 |
| Cingulum parahip. | .014 | .031 | .646 | .007 | .031 | .810 | -.003 | .031 | .920 | .006 | .031 | .841 | .005 | .031 | .878 |
| Inferior fronto-occipital | -.008 | .032 | .790 | -.021 | .032 | .516 | .002 | .032 | .944 | .014 | .031 | .661 | .011 | .031 | .729 |
| Uncinate | -.013 | .031 | .681 | -.015 | .031 | .631 | -.023 | .031 | .455 | -.030 | .031 | .323 | -.031 | .031 | .305 |
| Superior thalamic | -.013 | .032 | .672 | .005 | .032 | .866 | .016 | .032 | .606 | .021 | .031 | .500 | .021 | .031 | .513 |
| Anterior thalamic | .007 | .031 | .816 | .004 | .031 | .904 | .018 | .031 | .561 | .026 | .031 | .404 | .022 | .031 | .478 |
| Posterior thalamic | .011 | .031 | .725 | .009 | .031 | .762 | .028 | .031 | .365 | .037 | .031 | .236 | .035 | .031 | .263 |
| Corticospinal | -.045 | .031 | .144 | -.017 | .031 | .588 | -.023 | .031 | .459 | -.009 | .031 | .765 | -.008 | .031 | .805 |
| Forceps major | -.023 | .032 | .474 | -.045 | .032 | .151 | -.032 | .032 | .307 | -.020 | .031 | .531 | -.018 | .031 | .559 |
| Forceps minor | .017 | .032 | .591 | .007 | .031 | .833 | .024 | .031 | .451 | .016 | .031 | .605 | .014 | .031 | .661 |
| Acoustic | .024 | .031 | .449 | .001 | .031 | .982 | .006 | .031 | .847 | -.010 | .031 | .743 | -.015 | .031 | .638 |
| Medial lemniscus | .006 | .031 | .855 | .001 | .031 | .967 | .003 | .031 | .934 | -.005 | .031 | .867 | -.007 | .031 | .815 |
| Middle cerebellar | -.033 | .032 | .305 | -.025 | .032 | .440 | -.019 | .032 | .549 | .006 | .032 | .859 | .005 | .032 | .864 |

*Notes*. SCZ = schizophrenia; PRS = polygenic risk score; MDD = major depressive disorder; FA = fractional anisotropy. Reported results are for interactive effects between SCZ PRS and MDD status on tract FA, controlling covariates. No results were significant, even before correcting for multiple comparisons.

**Table S5 – Interactive effects between SCZ PRS and MDD status on MD by white matter tract.**

| | PRS threshold .01 | | | PRS threshold .05 | | | PRS threshold .10 | | | PRS threshold .50 | | | PRS all SNPs | | |
|---|---|---|---|---|---|---|---|---|---|---|---|---|---|---|---|
| | β | SE | p | β | SE | p | β | SE | p | β | SE | p | β | SE | p |
| Superior longitudinal | .010 | .030 | .753 | .022 | .030 | .459 | .010 | .030 | .734 | .014 | .030 | .638 | .017 | .030 | .563 |
| Inferior longitudinal | .028 | .030 | .354 | .041 | .030 | .168 | .028 | .030 | .345 | .027 | .030 | .370 | .032 | .030 | .285 |
| Cingulum cingulate | .025 | .029 | .398 | .032 | .029 | .266 | .022 | .029 | .451 | .030 | .029 | .292 | .032 | .029 | .268 |
| Cingulum parahip. | .000 | .032 | .989 | -.001 | .032 | .965 | .004 | .032 | .896 | -.009 | .032 | .773 | -.007 | .032 | .821 |
| Inferior fronto-occipital | .021 | .031 | .499 | .019 | .031 | .525 | .004 | .030 | .899 | .004 | .030 | .883 | .009 | .030 | .755 |
| Uncinate | .027 | .029 | .348 | .042 | .029 | .140 | .051 | .029 | .076 | .058 | .028 | .040 * | .062 | .028 | .029 * |
| Superior thalamic | -.004 | .027 | .875 | .014 | .027 | .603 | -.003 | .027 | .897 | -.010 | .027 | .708 | -.007 | .027 | .779 |
| Anterior thalamic | .012 | .028 | .672 | .031 | .028 | .274 | .018 | .028 | .526 | .017 | .028 | .535 | .021 | .028 | .456 |
| Posterior thalamic | -.012 | .029 | .678 | -.001 | .029 | .961 | -.020 | .029 | .485 | -.021 | .029 | .472 | -.017 | .029 | .550 |
| Corticospinal | .008 | .032 | .797 | .028 | .032 | .375 | .019 | .032 | .549 | .012 | .032 | .695 | .013 | .032 | .693 |
| Forceps major | .020 | .032 | .519 | .041 | .031 | .195 | .052 | .031 | .095 | .049 | .031 | .120 | .046 | .031 | .141 |
| Forceps minor | -.001 | .030 | .977 | .016 | .030 | .598 | .002 | .030 | .940 | .020 | .030 | .501 | .024 | .030 | .423 |
| Acoustic | .049 | .032 | .125 | .057 | .032 | .077 | .048 | .032 | .133 | .055 | .032 | .086 | .059 | .032 | .065 |
| Medial lemniscus | -.037 | .033 | .257 | -.013 | .033 | .697 | -.028 | .033 | .396 | -.008 | .032 | .813 | -.007 | .032 | .819 |
| Middle cerebellar | .001 | .032 | .979 | .017 | .032 | .589 | .023 | .032 | .482 | .033 | .032 | .306 | .033 | .032 | .298 |

*Notes*. SCZ = schizophrenia; PRS = polygenic risk score; MDD = major depressive disorder; MD = mean diffusivity. Reported results are for interactive effects between SCZ PRS and MDD status on tract MD, controlling covariates. * represents significant results at $p<.05$ before correcting for multiple comparisons. No results remained significant after FDR correction.
